# *Arabidopsis* Group I Pumilio RNA-binding factors are vital for embryo development and balancing between growth and stress resistance

**DOI:** 10.1101/2025.08.21.671196

**Authors:** Wenjuan Wu, Di Li, Danni Lin, Wangzhi Xu, Tianli Chen, Xiaomei Chen, Wei Guo, Zhengbiao Long, Xiang Xu, Xiaoyu Tu, Jirong Huang

**Author notes:** Correspondence: Jirong Huang. These authors contributed equally to this article (Wenjuan Wu, Di Li). The author responsible for distribution of materials integral to the findings presented in this article in accordance with the policy described in the Instructions for Authors (https://academic.oup.com/plphys/pages/General-Instructions) is Jirong Huang.

## Abstract

Pumilio (PUM) RNA binding proteins are crucial for regulating gene expression by binding to a conserved motif in the 3′-untranslated region (3′-UTR). Despite their importance, the role of PUM in plants is largely unexplored. Here, we investigated the functions of *Arabidopsis* group I PUMs (APUM1-APUM6), which are ubiquitously expressed and localized in the cytosol. While single *apum* mutants exhibit no visible phenotypes, CRISPR/Cas9-generated *apum1 apum2 apum3* triple mutants (*apum1/2/3*) display reduced growth in both vegetative and reproductive organs, alongside hypersensitivity to various stresses. Remarkably, *apum1/2/3/4* quadruple mutants are embryonically lethal, highlighting their essential role in embryo development. Transcriptomic profiling revealed that differentially expressed genes (DEGs) upregulated in *apum1/2/3* are enriched in pathways related to photosynthesis, stress responses and anthocyanin biosynthesis, while downregulated DEGs are associated with biotic stress defense and hydrogen peroxide metabolism. This suggests that APUM1/2/3 act as molecular hubs balancing plant growth and stress adaptation. Biochemical assays showed that recombinant APUM homologous domains bind to the 5’-UG**U**GUAUA-3’ core motif in the 3’-UTR of the transcription factor Production of Anthocyanin Pigment1 (PAP1), crucial for anthocyanin biosynthesis. Notably, single nucleotide substitutions, except for the third U, do not affect binding, while multiple mutations disrupt interaction. Consistently, *apum1/2/3* mutants accumulate significantly more anthocyanin than wild-type plants. Furthermore, we predicted 7053 potential target genes for APUM1/2/3, with 1609 present among the upregulated DEGs in *apum1/2/3*. Taken together, our findings demonstrate that group I APUMs are vital posttranscriptional regulators, providing a new perspective on the trade-offs between growth and stress resilience in plants.

## INTRODUCTION

The expression of eukaryotic genes is coordinately regulated by multiply levels including transcription, post-transcription, translation and post-translation. Post-transcriptional regulation is largely mediated by RNA binding proteins (RBPs) that influence the splicing, processing, transport, stability and translation of their target RNAs. One of the evolutionarily conserved RBPs in eukaryotes is the small Pumilio (PUM, meaning dwarf in Latin) family, which has been demonstrated to modulate a wide range of biological processes, such as development of the embryo and nervous system, maintenance of stem cells and germ cells, chemotactic cell movement and human diseases (Goldstrohm et al., 2018; Nishanth and Simon, 2020). Originally, *pum* was named for the *Drosophyla* mutants with less than the normal eight abdominal segments of embyos (Lehmann and Nüsslein-Volhard, 1987). Thereafter, *Drosophila* PUM and its *Caenorhabditis elegans* homolog, designated as the *fem-3* binding factor (FBF, regulating sexual fates in the hermaphrodite germ line), were almost simultaneously reported to be a new class of RNA-binding proteins, which specifically bind a conserved short motif in the 3′-untranslated region (3′-UTR) (Zamore et al., 1997; Zhang et al., 1997). To date, the PUM family proteins have been well characterized for the conserved PUM homologous domain (PUM-HD) or PUF (PUM/FBF) domain. Crystal structure analysis revealed that PUM-HD consists of eight PUM repeats and two pseudo-repeats at the N and C termini, which patch together to form a right-handed crescent structure, suggesting that the inner face of the domain binds to RNA while the outer face interacts with proteins (Wang et al., 2002). Each PUM repeat owns approximately 36 amino acid residues and form three consecutive α helices. The middle helix provides a binding interface with the target RNA (Muraro et al., 2008; Zhang et al., 1997). The three (12^th^, 13^th^ and 16^th^) conserved amino acid residues within each PUM repeat are responsible for binding to a single RNA base via hydrogen bonds, der Waals and base stacking interactions (Wang et al., 2002). Although different PUMs preferentially bind to RNA motifs of distinct length and sequences, the PUM recognition/response elements (PRE) share the 5′-UGUANAUA-3′ (N represents A, U, or C) motif, where the beginning 5′-UGU is the highly conserved core triplet for all PUM proteins (Lu and Hall, 2011). The best-documented mechanism of PUM-mediated repression of gene expression is through degradation of target mRNA by recruiting the Ccr4-Not deadenylase complex (CNOT) (Wharton et al., 1998; Wreden et al., 1997). Alternatively, PUMs can also repress protein translation through interfering with recognition of cap structure and attenuation of translational elongation (Friend et al., 2012). It has also been demonstrated that mammalian PUM1 and PUM2 can promote protein expression and enhance mRNA stability (Uyhazi et al., 2020). In summary, PUMs primarily act as a negative master regulator of post-transcriptional gene expression by deadenylating its target transcripts or inhibiting translation (Goldstrohm et al., 2007; Kedde et al., 2010).

Compared to yeast and mammalian where merely 6 and 2 PUMs are present, respectively, higher plants usually contain more PUM proteins. For example, there are 26, 20 and 19 PUMs in the *Arabidopsis*, rice and maize genomes, respectively, suggesting that they play a wide range of roles in post-transcriptional regulation of gene expression (Dedow and Bailey-Serres, 2019; Feng et al., 2023; Huh, 2021). To date, however, only a few of PUMs have been investigated in terms of biological functions and molecular mechanisms. The *Arabidopsis* 26 PUMs, named from *APUM1* to *APUM26,* are classified into five groups depending on the number of PUM repeats in PUM-HD (Huh, 2021). Bioinformatics analysis revealed that about 43% of the *Arabidopsis* gene transcripts contain the conserved APUM-binding sequence in the 3’-UTR regions, indicating that APUM proteins are possibly involved in regulating expression of a wide arrange of plant genes (Tam et al., 2010). The expression of *APUM* genes always display in a tissue- or organ-specific manner and is induced by environmental changes, while their subcellular localization have shown to be in the cytoplasm, nucleus and chloroplast (Abbasi et al., 2011). Interestingly, APUM5 is involved in suppressing infection of *cucumber mosaic*

*virus* by binding to viral RNA directly (Huh et al., 2013), in addition to participating in salt stress responses (Huh and Paek, 2014). The nucleolus-located APUM23 and APUM24 are required for rRNA processing, and functions in salt resistance and early embryo formation, respectively (Abbasi et al., 2010; Huang et al., 2014; Maekawa et al., 2018; Shanmugam et al., 2017; Zhang and Muench, 2015). APUM proteins were also detected in polysomal fractions during seed germination, indicating their involvement in active translation (Balarynová et al., 2025). Recently, APUM24 was reported to regulate seed maturation by reducing the mRNA stability of *BTB/POZMATH* (*BPM*) family genes, and overexpression of *APUM24* significantly improved seed yield (Huang et al., 2021). APUM9, whose expression is observed specifically during the seed maturation and imbibed seeds, negatively regulates the expression of ABA signaling-related genes during seed imbibition, promoting seed germination (Nyikó et al., 2019; Xiang et al., 2014). In addition, overexpression of APUM9 leads to abnormal leaf development, late flowering and enhanced heat tolerance, suggesting that APUM9 plays multiple roles in plant development and adaptation to environmental changes (Nyikó et al., 2019). The molecular mechanisms underlying APUM-regulated mRNA degradation have been shown to directly interact with decapping factor 2 (DCP2) of the mRNA 5’ capping complex (Nyikó et al., 2019) and carbon catabolite repressor 4 (CCR4) of the CCR4-NOT RNA deadenylase complex (Arae et al., 2019). Thus, plant pumilio proteins regulate gene expression through the same molecular mechanisms as those of yeast and animals.

Group I APUMs (from APUM1 to APUM6) were shown to share the highest similarity and identity in PUM-HD to those in *Drosophila* and human (Tam et al., 2010), suggesting that the biological function of group I APUMs is conserved and probably redundant in *Arabidopsis*. *APUM1*, *APUM2* and *APUM3* are arranged in tandem on chromosome 2, and have similarity higher than 85% (Francischini and Quaggio, 2009). It has been shown that PUM-HD of group I but not group II APUMs can bind to the same *NRE1* (*Nanos Response Element 1*) sequence as the target of *Drosophila* and human Pumilio 1, as well as to the 3’-UTR of *WUSCHEL*, *CLAVATA-1*, *PINHEAD/ZWILLE* and *FASCIA TA-2* transcripts in *Arabidopsis*, suggesting that they are involved in stem cell maintenance and differentiation (Francischini and Quaggio, 2009). However, no genetic and molecular biology evidence is available with regard to the role of Group I APUMs in plant growth and development. In this study, we investigated the biological significance of Group I APUM proteins, namely APUM1, APUM2 and APUM3 (APUM1/2/3), and found that APUM1/2/3 are functionally redundant and play an important role in plant grow, development and resistance to various stresses, suggesting that APUM1/2/3 coordinately regulate the balance between plant growth and development, and adaptation to environmental stresses.

## RESULTS

### Group I *APUMs* are ubiquitously expressed and function redundantly

To investigate the function of group I *APUM* genes, we first examined their expression patterns in various tissues of Arabidopsis plants. Quantitative PCR (qPCR) analysis showed that group I *APUM* genes were ubiquitously expressed in all examined tissues, but relatively at a lower level in stems, compared to those in rosette leaves under normal growth conditions (Figure 1A). In addition, different members of group I *APUMs* had their own expression patterns. For example, *APUM1* were expressed at a relatively higher levels in roots and flowers, while *APUM2* and *APUM3* at similar levels in all tissues (Figure 1A). Confocal microscopy assay showed that yellow fluorescence protein (YFP)-fused APUMs were all localized in the cytoplasm when transiently expressed in Arabidopsis protoplasts (Figure 1B), indicating that they play a role in the cytoplasm as shown in other species. Since the transcript of *APUM5* has been reported to be induced by biotic and abiotic stresses (Huh et al., 2013; Huh and Paek, 2014), we examined whether the expression of group I *APUMs* is induced by salt or osmotic stress. Total RNA was extracted from 10-day-old wild-type (WT) seedlings grown on the 1/2 MS medium and then transferred to the 1/2 MS liquid medium containing with or without 150 mM NaCl or 400 mM mannitol for 6 hours. qPCR analysis showed that compared to the control, salt stress significantly induced mRNA levels of all group I *APUMs* in a similar degree, whereas osmotic stress induced dramatic expression of from *APUM3* to *APUM6* but slight expression of *APUM1* and *APUM2* (Fig. 1C). These results suggest that APUM1 to APUM6 function either redundantly or differentially in plant responses to different stress. We then identified six T-DNA insertion single mutants (*apum1* to *apum6*) by PCR-based genotyping (Figure S1). As shown in Figure 1D, all single mutants did not exhibit any obvious phenotypes under normal growth conditions. Taken together, our results suggest that group I *APUMs* are ubiquitously expressed in plants and function redundantly, but individual *APUM* expression is largely dependent on environmental cues.

**Figure 1.**
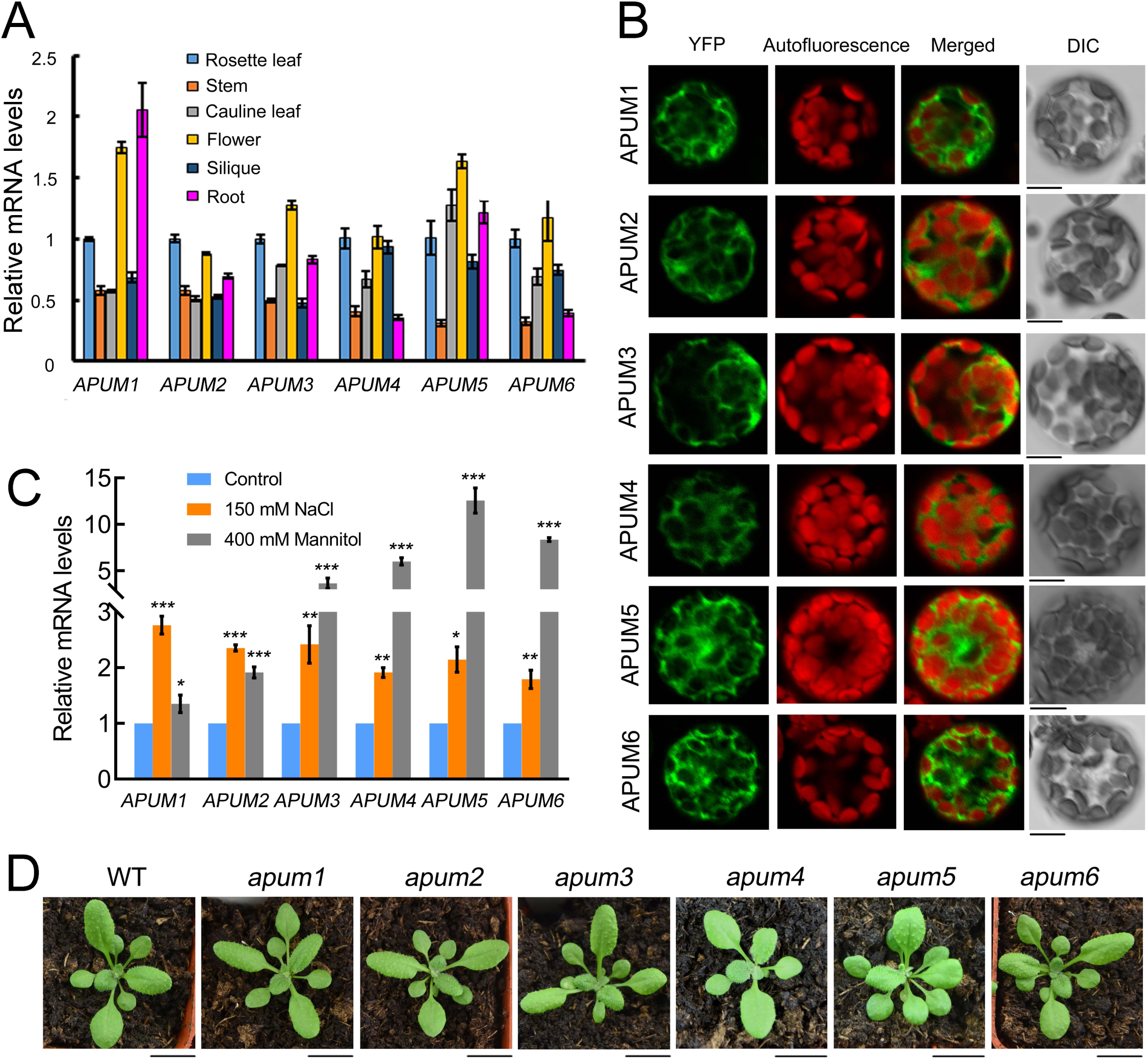
Characterization of group I APUM expression. **A)** Tissue specific expression patterns of *APUMs*. **B)** Subcellular localization of APUMs-GFP transiently expressed in *Arabidopsis* protoplasts. Bars, 10 μm. **C)** Relative expression levels of *APUMs* induced by salt and mannitol stresses. Expression levels of *APUMs* in seedlings without treatment are set to 1. *ACTIN2* was used as a reference to normalize expression of each set of the data. Data were shown means ± SD (n = 3). Significant differences were performed using Student’s *t*-test. *, *P* < 0.05; **, *P* < 0.01; ***, *P* < 0.001. Two biological replicates were analyzed with similar results. **D)** Phenotype of 20-day-old single *apum* plants grown in long day conditions. Bars, 1 cm.

### Group I APUMs are essential for plant survival and *apum1/2/3* triple mutants display pleiotropic phenotypes

To examine the role of *APUMs* in plant growth and development, we created *apum* multiple mutants. Considering that *APUM1* (*at2g29200*), *APUM2* (*at2g29190*) and *APUM3* (*at2g29140*) share the highest identity in group I *APUMs* and are strongly linked on chromosome 2 (Figure S2A), a single specific 20-nt targeting sequence within the guide RNA was designed to simultaneously knock out the three genes with CRISPR-Cas9 technology, and transformed into the wild type (WT) background. We identified two independent homozygous lines of the *apum1 apum2 apum3* triple mutant, named *apum1/2/3-13* and *apum1/2/3-17*. Both *apum1/2/3-13* and *apum1/2/3-17* mutants contain single base pair insertions in the target region of *APUM1*, *APUM2* and *APUM3* (Figure S2B), resulting in a premature termination of protein translation (Figure S2C). These results indicate that *APUM1*, *APUM2* and *APUM3* are simultaneously knocked out in the *apum1/2/3* triple mutant. We then crossed *apum1/2/3* with *apum4* to generate their quadruple mutant. Our results showed that the *apum1/2/3/4* quadruple mutant was embryo lethal (Figure 2A), and the silique of *apum1/2/3* was significantly shorter than that of WT (Figure 2B). These results suggest that group I APUMs are essential for plant survival.

**Figure 2.**
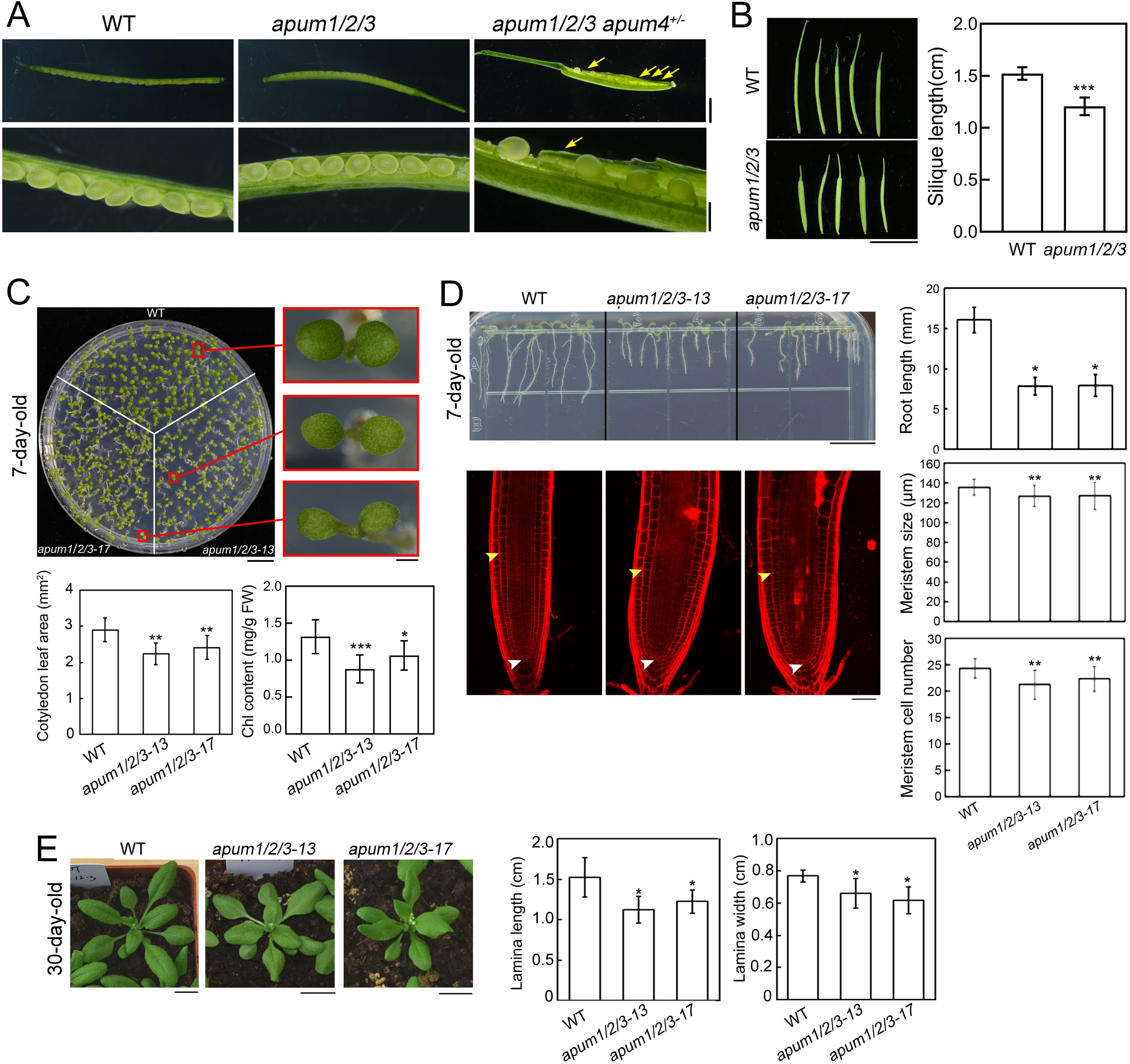
Functional analysis of *apum* multiple mutants. **A)** Yong seeds in siliques of WT, *apum1/2/3* and *apum1/2/3 apum4^+/-^* plants. Yellow arrowheads indicate the aborted embryos in *apum1/2/3 apum4^+/-^* siliques. Bars, 2 mm (up panel) and 0.5 mm (low panel). **B)** Phenotype (left panel) and length (right panel) of WT and *apum1/2/3* siliques at the same position of the main stem. Bars, 1cm. **C, D)** Phenotypes and quantification of 7-day-old WT and *apum1/2/3* seedlings grown on the 1/2 MS media in the long day condition. Bars in **C)**, 1 cm (left panel) and 0.5 mm (right panel). Bars in **D)**, 1 cm (up panel) and 50 μm (low panel). White and yellow arrowheads in the root stained with propidium iodide (PI) in **D)** indicate the quiescent center and the cell beginning to elongate, respectively. **E)** Phenotypes of 30-day-old WT and *apum1/2/3* plants grown in soil under long day conditions. Bars, 1 cm. The third pair of true leaves were quantified statistical analysis. All data were shown means ± SD (n ˃ 20). Statistical analysis was conducted with student’s *t* test. The significant differences between WT and mutants were indicated with stars. *, *P* < 0.05; **, *P* < 0.01; ***, *P* < 0.001.

Next, we characterized biological functions of group I APUMs using the *apum1/2/3* triple mutant. Phenotypic analyses showed that *apum1/2/3* displayed smaller and light green cotyledons in 7-day-old seedlings grown on half MS media containing 1% sucrose, compared to WT (Figure 1C). Consistently, *apum1/2/3* seedlings had significantly smaller cotyledons and less chlorophyll than WT (Figure 2C). In addition, *apum1/2/3* seedlings exhibited shorter roots than WT (Figure 2D). Quantification analysis showed that *apum1/2/3* had smaller root meristem size and less meristem cell number than WT (Figure 3D). When grown in soil, the leaf lamina of *apum1/2/3* mature plants (30-day-old) was smaller in both length and width than that of WT (Figure 2E). Thus, our data suggest that *APUM1/2/3* play a redundant role in regulating plant growth and development.

**Figure 3.**
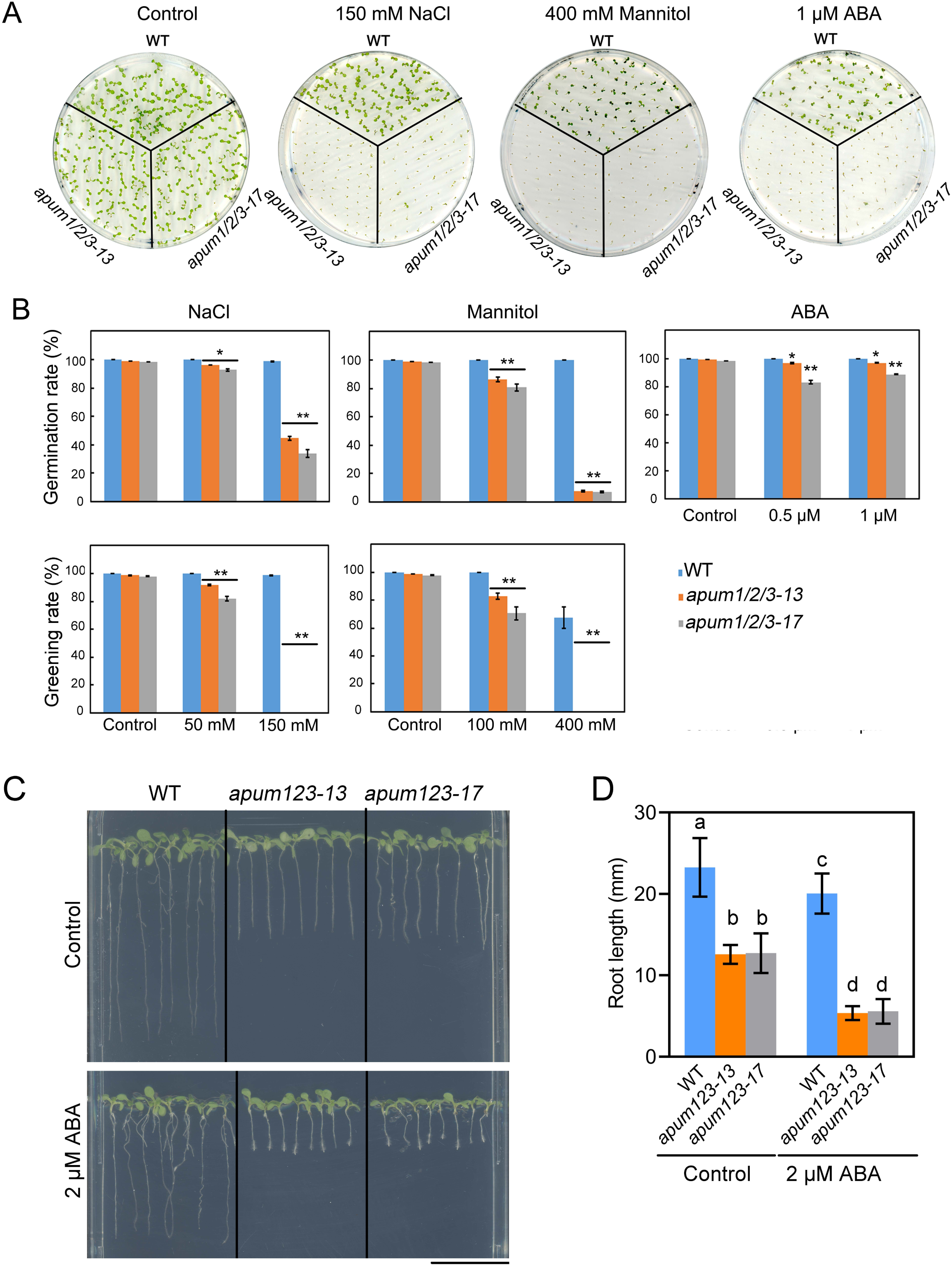
Hyposensitivity of *apum1/2/3* triple mutants to various stresses. **A)** Phenotypes of 7-day-old WT and *apum1/2/3* seedling treated with or without NaCl (150 mM), mannitol (400 mM) and ABA (1 μM ABA). **B)** Germination and greening rates of WT and *apum1/2/3* seeds grown on half MS media supplemented with indicated concentrations of NaCl, mannitol and ABA. Data were shown means ± SD (n = 3). Statistical analysis was conducted with student’s *t* test. Differences between WT and mutants were marked with star. *, *P* < 0.05; **, *P* < 0.01). **C)** Root sensitivity of WT and *apum1/2/3* to 2 μM ABA. Five-day-old seedlings grown on half MS media were transferred to the media with or without 2 μM ABA for another 4 days. Bars, 1cm. **D)** Quantification of root length in **C)**. Data were shown means ± SD (n > 30). Statistical analysis was conducted with one-way ANOVA. Different letters indicate significant differences (*P* < 0.05).

### *apum1/2/3* is hypersensitive to abiotic stress

Given that expression of group I *APUMs* is induced by abiotic stress, we investigated whether APUM1/2/3 are involved in plant resistance to environmental stress. To do this, WT and *apum1/2/3* seeds were germinated on the media containing different concentrations of NaCl or mannitol. Our results showed that only 34% and 7% of *apum1/2/3* seeds were germinated in 150 mM NaCl and 400 mM mannitol, respectively, a significant decrease in comparison with that of WT, whose germination rate remained almost constant with an increase in the concentration of NaCl or mannitol (Figure 3A, B). Likewise, the triple mutants displayed the same tendency in the rate of seedling greening as the germination rate (Figure 3A, B). However, almost no green seedlings were observed when *apum1/2/3* seeds were germinated on the media with 150 mM of NaCl or 400 mM of mannitol (Figure 3B). In addition, we tested the sensitivity of *apum1/2/3* mutants to ABA. Our data showed that both 0.5 and 1 μM treatments significantly reduced the germination rate of *apum1/2/3* seeds but not of WT (Figure 3A, B). The cotyledon greening of *apum1/2/3* mutants was completely blocked by 1 μM ABA, which had no effect on WT’s greening (Figure 3A). To further analyze the sensitivity of the triple mutant to ABA, we germinated seeds of WT and *apum1/2/3* on 1/2 MS media for five days, and then transferred seedlings to new 1/2 MS media containing with or without 2 μM ABA and cultured for another four days. Our results showed that the root length of *apum1/2/3* is shorter than that of WT in the absence of ABA, while become much shorter in the presence of ABA (Figure 3C, D). These results indicate that *APUM1/2/3* play an important role in plant adaptation to abiotic stress.

### PUM-HDs of APUM1/2/3 bind to the *NRE* sequence of *Drosophila* Pumilio

Previous studies reported that PUM-HD of group I APUMs can bind to the *NRE* target of *Drosophila* Pumilio using the yeast three-hybrid system (Francischini and Quaggio, 2009). To verify whether PUM-HDs of APUM1/2/3 bind to the NRE target, we purified GST-fused recombinant APUM1-HD, APUM2-HD and APUM3-HD proteins (Figure 4A). RNA electrophoresis mobility shift assay (REMSA) showed that all three PUM-HDs but not the GST control were able to bind the *NRE* sequence (Figure S3). We then used APUM1-HD to further characterize binding feature to the *NRE* sequence. Our results indicated that APUM1-HD interacted with the CY5-labeled NRE probe (P1) in a dose-dependent manner, and their interaction was disrupted by the unlabeled probe (C1) (Figure 4B). Notably, a single mutation of G to U in the NRE probe (P2) led to no interaction between P2 and APUM1-HD (Figure 4B), indicating that the second ‘G’ in the PRE core ‘UGUAUAUA’ is critical for binding. As shown in Figure 4C, APUM3-HD were detected the same binding characteristic to *NRE* as APUM1-HD. Taken together, our data demonstrate that APUM1/2/3-HDs can specifically bind to the NRE element.

**Figure 4.**
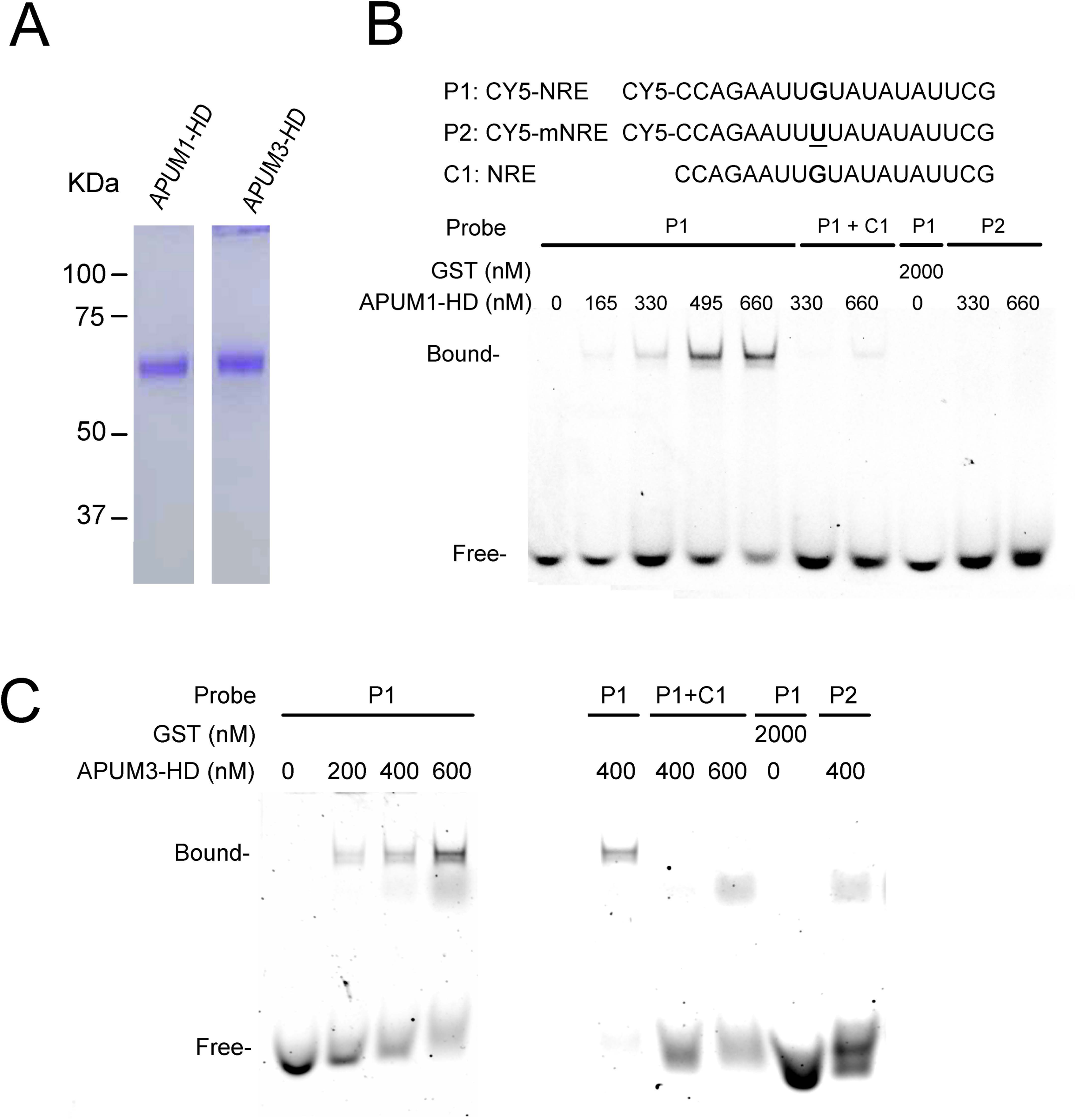
Binding assays between APUM homologous domain (HD) and the NRE (Nanos Response Element) motif. **A)** Purified recombinant APUM1-HD and APUM3-HD proteins from *E. Coli*. The gel was stained with Coomassie brilliant blue. **B, C)** REMSA analysis of the binding of APUM1-HD in **B)** and APUM3-HD in **C)** with NRE targets. P1, Cy5-labeled NRE probe; P2, Cy5-labeled mutated NRE probe; C1, unlabeled NRE probe. Bound represents the protein binding to Cy5 probe. Free represents the unbound Cy5 probe.

### APUM1/2/3 regulate expression of genes involved in photosynthesis, stress response and anthocyanin biosynthesis

To elucidate mechanisms underlying APUM1/2/3-regulated plant pleiotropic phenotypes, we profiled transcriptomes of WT and *apum1/2/3* seedlings grown under normal conditions. Principal component analysis (PCA) showed that transcriptomic data from the six samples (each genotype with three biological replicates) were distinctly clustered into WT and mutant groups, indicating of both high reproducibility and significant transcriptional differences between the genotypes (Figure 5A). Global gene expression analysis showed that there were a total of 8306 differentially expressed genes (DEGs, *p* ˂ 0.05) in *apum1/2/3*, compared to those in WT. Among these DEGs, 4249 (405 with fold changes > 2) were upregulated while 4057 (954 with fold changes > 2) were down-regulated (Figure 5B, Table S1-S3). To finely map the biological processes that were significantly altered by *APUM1/2/3* mutations, we selected DEGs with more than 2-fold changes either up- or down-regulated. Gene ontology (GO) enrichment analysis showed that the up-regulated DEGs were extensively enriched in photosynthesis including light reaction, light harvesting and pigment biosynthesis, response to light intensity and Jasmonic acid, and anthocyanin biosynthesis (Figure 5C, Table S2). For example, *APUM1/2/3* mutations led to a significant increase in expression of *Sig2* (*Sigma 2*, a transcription factor for the plastid encoded polymerase), *CSK* (*Chloroplast sensor kinase* encoding a two-component histidine kinase linking the plastoquinone redox signal with photosystem gene expression), *LHC* genes (*LHCB1*, *LHCB2*, *LHCB3*, *LHCB6*, *LHCA2* and *LHCA4*), *DEG5* encoding ATP-independent protease involved in the photodamaged D1 protein degradation of photosystem II, *STN7* encoding chloroplast thylakoid–associated serine-threonine protein kinase required for the phosphorylation of the major LHCII and for state transition, *BPG3* encoding BRZ-insensitive-pale green 3 required for the greening of leaves and related to brassinosteroid signaling, etc.

**Figure 5.**
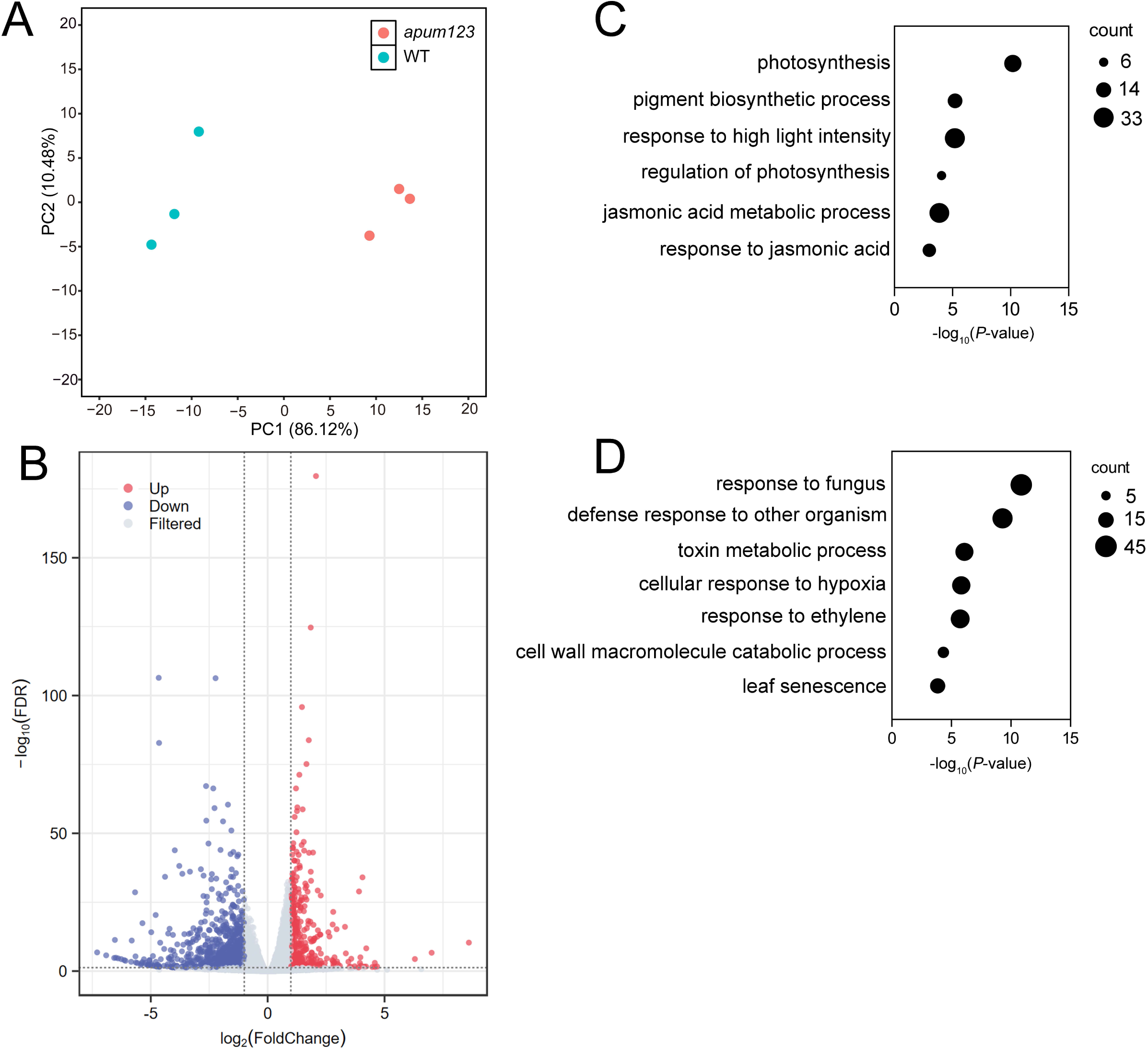
Effect of *APUM1/2/3* mutation on global gene expression. **A)** PCA analysis of the three independent transcriptomic data from WT and *apum1/2/3*. **B)** Volcano map of differentially expressed genes (DEGs) in *apum1/2/3*. Significantly upregulated and downregulated DEGs are shown as red and blue dots, respectively, while genes without significant difference between WT and *apum1/2/3* are shown as gray dots. **C, D)** GO enrichment analysis with significantly upregulated in **C)** or downregulated in **D)** DEGs (fold changes ˃ 2) in *apum1/2/3*.

Consistently, there are 4 DEGs encoding enzymes for chlorophyll biosynthesis, such as CAO (chlorophyll *a* oxygenase), GluTR (glutamyl-tRNA reductase), CHLH/GUN5 (H subunit of magnesium chelatase/genomes uncoupled 5, which also plays a key role in plastid-to-nucleus signal transduction), and DVR (3, 8-divinyl protochlorophyllide *a* 8-vinyl reductase). In addition, several genes enriched in pigment biosynthesis were involved in anthocyanin biosynthesis and regulation, such as *PAP1*, *TT3, TT18* and *UF3GT*, which encode production of anthocyanin pigment 1 (acting as a key transcription factor of anthocyanin biosynthesis), dihydroflavonol reductase, anthocyanindin synthase, and UDP-glucose:flavonoid 3-*O*-glucosyltransferase, respectively. In JA biosynthesis, we found that the upregulated genes in *apum1/2/3* include *LOX3* and *AOS*, which encode 13-lipoxygenase 3 and allene oxide synthase, respectively, and catalyze the first two steps in the octadecanoid pathway for JA biosynthesis, while *JMT* encoding jasmonic acid carboxyl methyltransferase catalyze formation of methyl jasmonate from JA. These data suggest that APUM1/2/3 play an important role in coordinately regulating photosynthesis, particularly in the process of light reaction, and environmental adaptation.

On the other hand, the down-regulated DEGs were mainly enriched in defense responses to biotic stress and the hydrogen peroxide metabolic process (Figure 5D, Table S3). For instance, expression of a number of genes encoding for receptor-like kinases such as FRK1/SIRK (FLG22-induced receptor-like kinase 1), CRK4 (cysteine-rich RECEPTOR-like kinase 4), CRK5, CRK6, CRK20 and CRK32, which is involved in early defense signaling to pathogen infection, were decreased in *apum1/2/3*. We also found that a series of the downstream genes involved in pathogen-associated responses, such as *SBT3.3* (subtilase 3.3 function as a receptor located in the plasma membrane activating downstream immune signaling processes), *WRKY51* (WRKY DNA-binding protein involved in JA inducible defense responses), *FMO1* (flavin-dependent monooxygenase 1, promoting resistance and cell death at pathogen infection sites), and *PR4* (pathogenesis-related 4, binding to the antifungal chitin) were downregulated in *apum1/2/3*. It was noteworthy that *APUM1/2/3* mutations resulted in suppression of salicylic acid (SA)-dependent defense-related gene expression. For example, *SARD1* encoding systemic acquired resistance deficient 1 acts as a key regulator for *ICS1* (isochorismate synthase 1) induction and SA synthesis; *WRKY70* functions as activator of SA-dependent defense genes and a repressor of JA-regulated genes; and *AZI1* (azelaic acid induced 1) and *EARLI1* (early *Arabidopsis* aluminum induced 1, a putative lipid transfer protein) are involved in priming the SA induction and systemic immunity triggered by pathogen SAR). Interestingly, there were 17 class III peroxidase genes (*PRX*) in the downregulated DEGs including *PRX01, PRX4, PRX33, PRX17, PRX37, PRX62* and *PRX56*, which target to cell wall and are involved in lignin polymerization and redox homeostasis. Thus, our data suggest that *APUM1/2/3* are also required for plant defense to biotic and abiotic stresses.

In summary, results derived from transcriptomic analysis are consistent with the phenotypes of the *apum1/2/3* mutants, and suggest that APUM1/2/3 play an important role in coordinately regulating plant photosynthesis and adaptation to environmental stresses.

### Identification of APUM-HD binding core motifs in plants

To identify candidate genes directly targeted by APUM1/2/3, we first searched the 3’-UTR regions of the genes upregulated more than 1.5-fold in *apum1/2/3* for the NRE core motif 5’-UGUAUAUA-3’ or its variants with one-altered nucleotide due to the second essential G nucleotide (Figure 4) and no effect of any other mutations on binding (Figure 6). Our results revealed 37 genes that contain at least one predicted APUM-binding sites in the 3’-UTR (Table S4). Interestingly, we found that a number of important APUM1/2/3 targets were involved in GO enriched processes, such as *PAP1* (encoding a bHLH transcription factor for anthocyanin biosynthesis), *JAZ13* (a repressor of JA signaling), *bHLH38* (in the regulation of osmotic stress response and iron homeostasis), and *LHCB1* (in light harvesting for photosynthesis).

**Figure 6.**
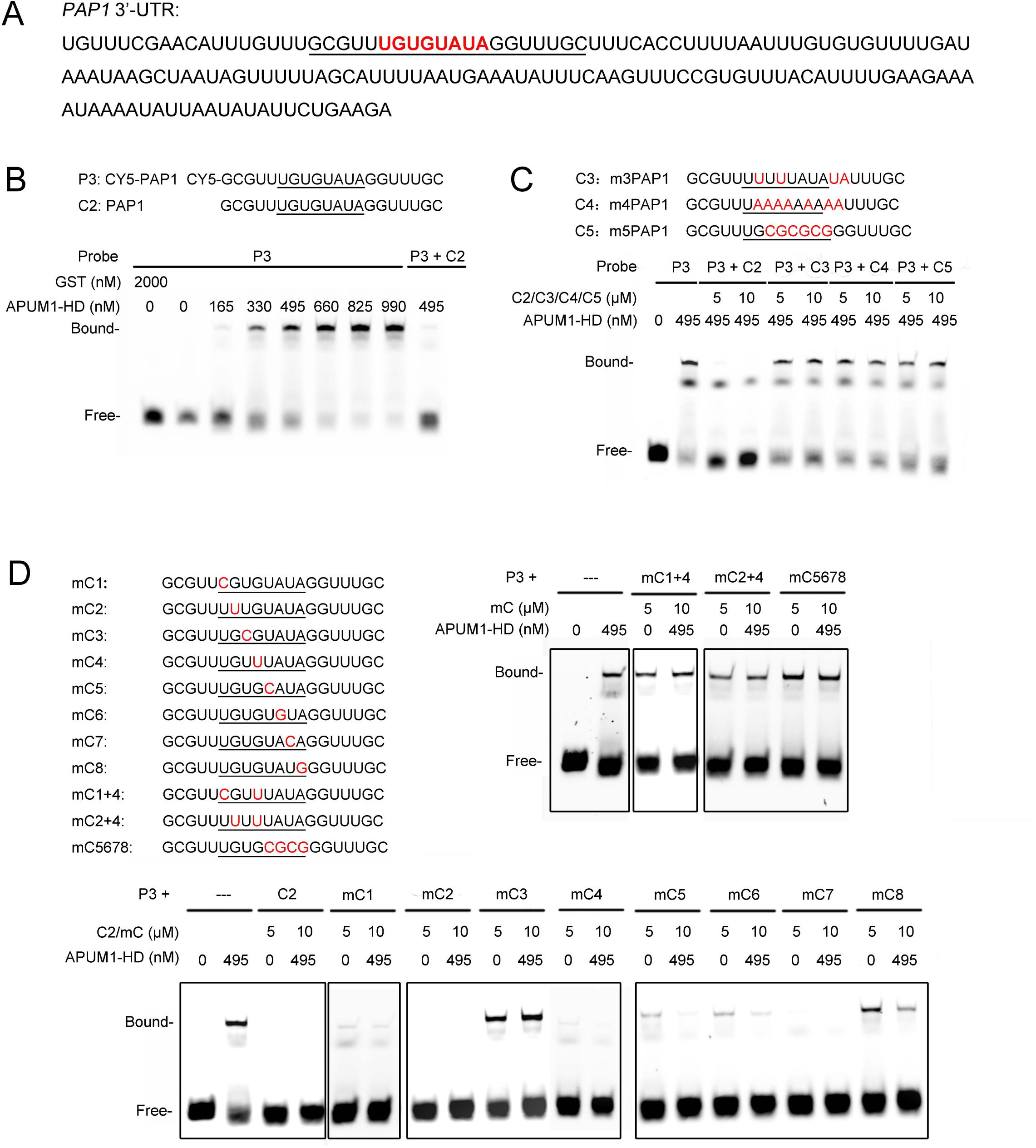
Finely mapping the key nucleotides in the 3’-UTR of *PAP1* mRNA binding to APUM1-HD. **A)** The 3’-UTR sequence of *PAP1* mRNA. Underlined nucleotides were used for making CY5-labelled P3 and unlabeled C2 probes in **B)**. The core binding motif 5’-UGUGUAUA-3’ is highlighted with red. **B)** REMSA analysis of APUM1-HD binding to the *PAP1* probe. **C)** REMSA analysis of APUM1-HD binding to mutated *PAP1* probes C3, C4, and C5 without any label. **D)** REMSA assays of APUM1-HD binding to mutated unlabeled probes shown in left panel. Bound represents the protein binding to Cy5 labeled probes. Free represents the unbound Cy5 labeled probe.

To confirm whether APUM1/2/3 can bind to the predicted RNA motif *in vitro*, we selected a representative one, 5’-UGUGUAUA-3’, in the 3’-UTR of *PAP1* (Figure 6A). To test the interaction between APUMs and the 3’-UTR of *PAP1*, we purified recombinant APUM1-HD from bacteria and prepared for CY5-labeled (P3) or unlabeled probes (C2) containing the 5’-UGUGUAUA-3’ motif. REMSA results showed that APUM1-HD could bind to the probe (P3) in a dose-dependent manner, while unlabeled competitive probe C2 completely competed the binding between APUM1-HD and P3 (Figure 6B). In addition, we found that APUM1-HD failed to bind C3, C4 and C5 unlabeled probes, where more than two nucleotides were mutated in the core sequence or plus its flanking sequence (GG) (Figure 6C). These results indicate that APUM1-HD specifically binds to the 3’-UTR of *PAP1* mRNA through a core binding motif. Subsequently, single-nucleotide substitutions were introduced into the *PAP1* core sequence 5’-UGUGUAUA-3’. Our data demonstrated that except for the third uridine (U) substitution with cytosine (C), other single nucleotide mutations could still compete for the interaction of CY5-labeled *PAP1* (P3) with APUM1-HD (Figure 6D), suggesting that the third uridine in 5’-UG**U**GUAUA-3’ plays a key role in binding to APUM-HD. Consistently, if two or more nucleotides of the other seven nucleotides were simultaneously mutated, labelled as mC1+4, mC2+4 and mC5678 in Figure 6D, their bindings to APUM1-HD were significantly reduced or even loss (Figure 6D). Taken together, our data indicate that APUM1 specifically bind to the core motif at the 3’-UTR region, and the binding can tolerate one nucleotide substitution except for the third uridine in 5’-UG**U**GUAUA-3’.

Based on these rules, we predicted genome-wide APUM-binding sites in *Arabidopsis* mRNA 3’-UTR regions. Our results showed a total of 7053 genes that could be potentially regulated by APUM1/2/3 (Table S5). Interestingly, 37.88% (1609) of the 4249 upregulated DEGs in *apum1/2/3* were present in the predicted genes (Table S5). These results indicate that group 1 APUMs regulate expression of genes in a whole genomic level.

### Mutations in *APUM1/2/3* lead to anthocyanin overaccumulation

Given that *PAP1* expression is regulated by APUM1/2/3, we investigated the role of *APUM1/2/3* in anthocyanin biosynthesis. To do this, we cultured WT and *apum1/2/3* seeds in 1/2 MS liquid media without sucrose for 4 days, and then transferred seedlings to the media containing with or without 5% sucrose for another 2 days to induce anthocyanin synthesis. The results showed that both *apum1/2/3* and WT seedlings did not accumulate anthocyanin in the absence of sucrose, whereas *apum1/2/3* seedlings accumulated more anthocyanin in the cotyledon and hypocotyl than WT in the presence of 5% sucrose (Figure 7A). Quantification analysis indicated that *apum1/2/3* accumulated 1.6-fold anthocyanin of WT in the presence of 5% sucrose (Figure 7B). qPCR analysis showed that after sucrose induction, expression levels of representative key genes for the anthocyanin synthesis pathway and transcription regulatory factors, such as *PAP1* (*production of anthocyanin pigment 1*), *TT8* (*TRANSPARENT TESTA 8*), *AT5MAT* (anthocyanin *5-O-glucoside-6″-O-malonyltransferase*), *UF3GT* (*UDP-glucose/flavonoid 3-O-glucosyltransferase*), and *DFR* (*dihydroflavonol 4-reductase*), were significantly up-regulated in *apum1/2/3*, compared to WT (Figure 7C). Thus, our results suggest that *PAP1* are directly regulated by APUM1/2/3.

**Figure 7.**
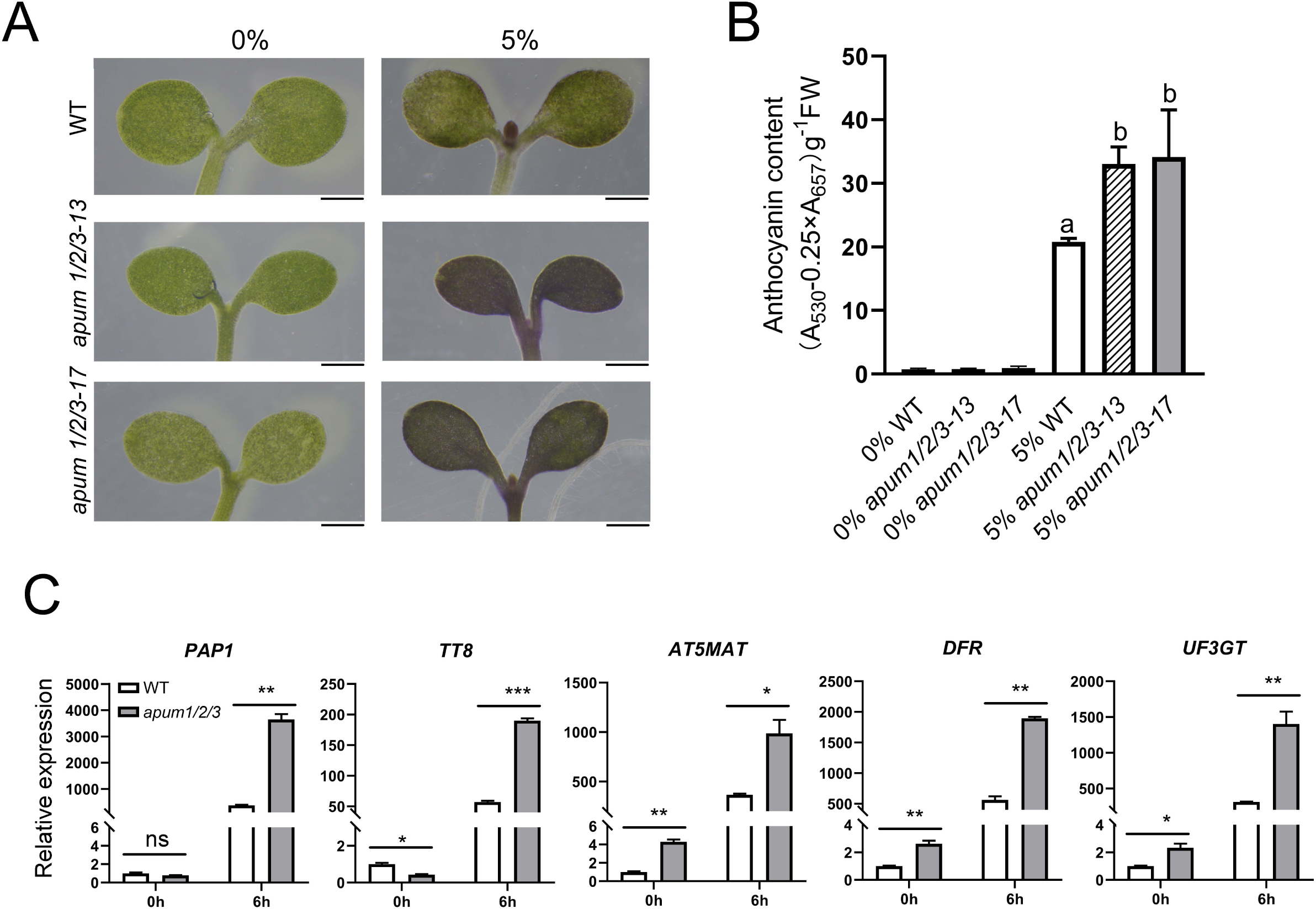
Mutations in *APUM1/2/3* lead to overaccumulation of anthocyanin. **A)** *apum1/2/3* cotyledons produced more anthocyanin than WT. Four-day-old seedlings cultured in 1/2 MS liquid media without sugar were transferred to the media with or without 5% of sucrose for additional 2 days. Bars, 0.5 mm. **B)** Anthocyanin content of the seedlings shown in **A)**. Data were mean ± SD (n = 3). Statistical analysis was conducted with one-way ANOVA. Different letters indicate significant difference among genotypes. **C)** Quantitative real-time PCR analysis of expression levels of genes encoding regulators and enzymes of anthocyanin biosynthesis. Total RNA was extracted from seedlings sampled at 0 h and 6 h of sucrose treatment. Data were shown means ± SD (n = 3). Expression levels of the genes in WT seedlings without sucrose treatment are set to 1. *ACTIN2* was used as a reference to normalize expression of each replicate. Statistical analysis was performed with student’s *t* test. Stars indicates significant difference between WT and *apum1/2/3* mutants. *, *P* < 0.05; **, *P* < 0.01. Two biological replicates were analyzed with similar results.

## DISCUSSION

In this study, we characterized the basic functions of group 1 APUMs, mainly APUM1/2/3, through genetic, physiological, biochemical and transcriptomic approaches. We found that the triple *apum1/2/3* mutant exhibits a slower growth rate in both leaves and roots, and hypersensitive to environmental stresses including salt and osmosis, compared to WT. These morphological and physiological phenotypes are strongly supported by the transcriptomic data showing that APUM1/2/3 coordinately regulate expression of genes involved in the biological processes including photosynthesis, pathogen infection, and secondary metabolite biosynthesis. Further analysis revealed that APUM1/2/3 negatively control anthocyanin synthesis via direct binding to the core motif of 5’-UGUGUAUA-3’ in the 3’-UTR of *PAP1*, which encodes the key transcription factor for anthocyanin synthesis under various stresses. Notably, we found that the *apum1/2/3/4* quatuble mutant is embryoic lethality. Thus, we conclude that group 1 APUMs are essential for plant growth, and APUM1/2/3 play an important role in plant growth and adaptation to environmental stresses via specifically binding to the 3’-UTR of their target mRNAs.

### Overlapping functions of APUM1/2/3 in plant growth

The six APUMs in group I share the approximately 50% identity and 75% similarity to the *Drosophila* PUM-HD domain, which is the highest one among the six groups (Francischini and Quaggio, 2009). *APUM1* (At2g29200), *APUM2* (At2g29190), and *APUM3* (At2g29140) form a tightly clustered group on chromosome 2 with more than 85% amino acid sequence identity, suggesting recent gene duplication events and potential functional overlap (Arae et al., 2019). This raises the question whether these APUMs are functionally redundant. Our analysis of gene expression patterns showed that *APUM1-APUM6* are ubiquitously expressed in all examined tissues and their products were all localized in the cytoplasm, which supports that these isoforms may compensate for one another in biological contexts. Consistently, our genetic data showed that the six single mutants, from *apum1* to *apum6*, exhibit no defective phenotype in plant growth, while quatuble *apum1/2/3/4* and triple *apum1/2/3* mutants display the embryo lethal and pleiotropic phenotypes, respectively. For example, *apum1/2/3* shows a slow growth rate of plant leaves and roots, and hypersensitivity to environmental stimuli. Further analysis indicated that short root length of *apum1/2/3* was due to the smaller size of root meristems. These results are in agreement with the proposal that group I APUMs (from APUM1 to APUM6) share overlapping involvement in shoot stem cell maintenance, potentially targeting common pathways like WUSCHEL regulation (Huh, 2021). In fact, APUM2 has been reported to bind transcripts involved in shoot meristem maintenance (Francischini and Quaggio, 2009), and the near-identical APUM1 and APUM3 might share this capacity, which ensures robustness in stem cell regulation. The functional redundancy of group I APUMs is further supported by their shared localization to cytoplasmic processing bodies (P-bodies), where they interact with CCR4-NOT deadenylase complexes to regulate mRNA stability (Arae et al., 2019). On the other hand, functional specialization is evident from distinct roles reported for individual APUM members (Abbasi et al., 2010; Huang et al., 2021; Huang et al., 2014; Huh and Paek, 2014; Maekawa et al., 2018; Nyikó et al., 2019; Shanmugam et al., 2017; Un Huh and Paek, 2013). These distinct phenotypes and mechanisms highlight non-redundant roles tied to unique domains, subcellular localizations, and tissue-specific expression patterns. For example, APUM9 expression is restricted to the seeds and dehiscence zone of siliques due to a transposable element (TE) insertion in the promoter (Nyikó et al., 2019), and APUM9 expression is strongly correlated with changes in seed dormancy level (Xiang et al., 2014). It is possible that the evolutionary expansion of group 1 APUMs in *Arabidopsis* has led to functional specialization, driven by divergent domains, expression dynamics, and target RNA specificities, although their primary functions overlap. It will be interesting to address whether APUM1/2/3 can function in a specialized manner.

### APUM1/2/3 coordinately regulate various biological processes

Since plants are sessile organisms, it is particularly important to quickly respond and adapt to environmental changes. Post transcriptional regulation has been recognized as a rapid and effective mechanism to protect plants from damage caused by various biotic and abiotic stresses, in which RNA binding proteins (RBPs) play an essential role in turning on/off expression of a specific set of genes (Moore, 2005). APUM proteins modulate mRNA stability, localization, or translation efficiency, thereby fine-tuning gene expression critical for diverse biological processes. Photosynthesis, one of the important processes for plant growth and development, is vulnerable to be influenced by environmental cues. However, it remains totally unclear whether plant photosynthesis is regulated by APUM proteins. In this study, GO enrichment analysis of the transcriptome revealed that the upregulated DEGs identified in *apum1/2/3* were significantly involved in the light reaction process of photosynthesis including the synthesis of photosynthetic pigments and their binding proteins (LHCA and LHCB), and in the regulatory network to maintain homeostasis of photosynthesis. For example, *SIG2* and *CSK* play an important role in regulating expression of the plastid-encoded photosynthetic genes, *STN7* is required for state transition that redistributes energy between PSI and PSII to avoid over-excitation under stress, and *DEG5* is involved in photodamaged D1 degradation in the PSII repair. It is suggested that PSII is over-excited in the *apum1/2/3* mutant, producing a higher level of reactive oxidative species (ROS) including superoxide anion (O₂⁻•), hydrogen peroxide (H₂O₂) and hydroxyl radical (•OH) in the mutant chloroplasts. These ROS can act as a signal recognized by various sensors to trigger the retrograde signaling from chloroplasts to the nucleus or stimulate biosynthesis of phytohormonal precursors for JA and ABA in chloroplasts. Consistently, we found that *GUN5* that plays an important role in retrograde signaling and three key genes (*LOX3, AOS* and *JMT*) in JA biosynthesis were upregulated, indicating that APUM1/2/3 are involved in amplifying systemic defense signaling and defending against herbivores and necrotrophic pathogens while modulating photosynthetic acclimation in the mutant. However, we found that JAZ13, a non-TIFY repressor of JA signaling, was significantly upregulated. JAZ13 was reported to interact with MYC2 and the co-repressor TOPLESS but not with NINJA and other JAZs, and overexpression of JAZ13 results in reduced sensitivity to JA, attenuation of wound-induced expression of JA-response genes, and decreased resistance to insect herbivory (Thireault et al., 2015). Thus, upregulated expression of JAZ13 may allow fine-tuning of JA-mediated transcriptional responses. In addition, upregulated expression of *BPG3* also revealed that APUM1/2/3 play a role in BR-regulated chloroplast development.

We cannot exclude the possibility that ROS acts as a destroyer of biomolecules to inhibit plant growth since the mutant displays a smaller plant body in comparison with WT. Unexpectedly, our transcriptomic analysis did not show that *APUM1/2/3* mutations have a significant effect on expression of cell cycle-related genes, which have been reported to be regulated by group I APUMs in plants and animals (Francischini and Quaggio, 2009; Lin et al., 2019; Racher and Hansen, 2012). One possibility could be due to the presence of other members (APUM4/5/6) of group I APUMs in the triple mutant. Collectively, our data indicate that APUM1/2/3 are critical modulators that balance photosynthetic activity with stress resilience by facilitating transcriptional reprogramming.

Given that APUMs negatively regulate gene expression via direct binding to target mRNAs, we thus propose that the downregulated DEGs are the secondary effects of *APUM1/2/3* mutations. GO analysis of the downregulated DEGs were over-presented in responses to bacteria and ROS (Figure 5D). *APUM1/2/3* mutations result in reduced expression of genes not only for the SA biosynthetic pathway (such as *SARD1* and WRKY70) but also for SA priming and systemic immunity (AZI1 and EARL1). In addition, many genes encoding class III peroxidases that involve in lignin polymerization and redox balance were also downregulated in *apum1/2/3*. This phenomenon could be attributed to antagonistic action between JA and SA. Although the mechanism by which SA suppresses expression of JA-responsive genes is well documented, the pathway for inhibition of SA responsive genes by JA remains unclear (Roychowdhury et al., 2024). Hou and Tsuda (2022) reported that the JA-mediated suppression of SA signaling was via decreasing SA biosynthesis and accumulation rather than hampering SA signaling (Hou and Tsuda, 2022). JA can trigger the expression of the C2H2 zinc finger transcription factor ZAT18 via MYC2, which suppresses the expression of *ICS1* and *EDS1* (*Enhanced Disease Susceptibility 1*), thereby decreasing accumulation of SA (Gao et al., 2022). Indeed, we observed that *ICS1* and many SA-response genes are downregulated in the mutant. Normally, SA suppresses JA signaling via NPR1-WRKY70-mediated repression of JA-responsive genes such as MYC2 (Li et al., 2004; Nomoto et al., 2021). In contrast, in the mutants, downregulation of *WRKY70* removes its repression of JA pathways, yet JA-responsive genes like WRKY51 (a positive regulator of JA defenses) are also suppressed. This dual SA-JA dysregulation suggests APUM1/2/3 act as hubs for integrating stress signals, ensuring transcriptional activation of defense genes while balancing antagonistic hormone pathways. In addition, *APUM1/2/3* mutations result in decreased expression of a set of genes involved in the early defense signaling such as FRK1/SIRK and CRKs. It is worthy to test whether *apum1/2/3* mutants are hypersensitive to biotrophic and necrotrophic pathogens, which are primarily regulated by SA and JA, respectively. Taken together, our data highlight that APUM1/2/3 act as a hub to integrate photosynthetic efficiency with stress adaptation. The molecular interplay involves retrograde signaling, hormonal crosstalk, and redox regulation, offering targets for engineering crops with improved stress resilience without compromising productivity.

### Posttranscriptional regulation of anthocyanin biosynthesis by APUM1/2/3

Anthocyanin accumulation has been recognized as a well-known biomarker when plants suffer from environmental stresses, such as high light, UV radiation, drought, cold, nutrient deficiency, and pathogen attack. These pigments act as antioxidants to scavenge reactive oxygen species (ROS) and protect macro-molecules from UV damage. Anthocyanin biosynthetic genes are specifically regulated by the ternary MYB-bHLH-WD40 (MBW) complex, which is composed of the R2R3 MYB (MYB) and bHLH transcription factors and WD40-repeat protein (Gonzalez et al., 2008; Ramsay and Glover, 2005). Since the MYB subunit, which is also called production of anthocyanin pigment (PAPs), is induced by multiple stimuli while bHLH and WD40 subunits are constitutively expressed, the MYB subunit plays a primary role in initiating anthocyanin biosynthesis. Contrast to the well-characterized MBW-mediated transcriptional regulation of anthocyanin biosynthesis (Gonzalez et al., 2008; Ramsay and Glover, 2005), its posttranscriptional mechanisms remain largely unknown. mRNA degradation is an essential process for posttranscriptional regulation of gene expression in eukaryotes. To date, two general pathways for mRNA turnover have been identified in eukaryotic cells. One is the deadenylation-dependent decay, by which Poly(A) tails are first shortened by deadenylases such as CCR4-NOT complex and then mRNAs are decapped by DCP1/DCP2 complexes, exposing the transcript to digestion by exonuclease, another is the endonuclease-dependent cleavage either by sequence-specific endonucleases or in response to miRNAs or siRNAs (Parker and Song, 2004). Several microRNAs have been reported to regulate anthocyanin biosynthesis via directly or indirectly targeting transcripts encoding MYB transcription factors. For example, Hsieh *et al*. (2009) demonstrated that miR828 induces cleavage of the *Trans-Acting SiRNA Gene 4* (*TAS4*) mRNA, generating small interfering RNAs (*ta-siRNAs*) that specifically target *PAP1*, *PAP2* and *MYB113* in Arabidopsis (Hsieh et al., 2009). Similarly, miRNA858 directly target anthocyanin repressor *MYBL2*, which negatively regulate anthocyanin accumulation by inhibiting the MYB subunit to form the MBW transcriptional activation complex (Wang et al., 2016). Conversely, miRNA156 promotes anthocyanin biosynthesis in *Arabidopsis* by destabilizing *Squamosa Promoter binding-Like* (*SPL*), which repressed formation of the MBW complex (Gou et al., 2011). However, it remains unclear whether deadenylation-dependent RNA degradation is involved in anthocyanin biosynthesis. In this study, we found that the *apum1/2/3* seedling is vulnerable to accumulate anthocyanin. In agreement with this phenotype, transcriptional analysis showed that *APUM1/2/3* mutations lead to a significant increase in the expression level of examined genes involved in anthocyanin biosynthesis and regulation (Figure 7). Bioinformatics analysis indicated that there is an APUM-binding core sequence 5’-UGUAUAUA-3’ in the 3’-UTR of *PAP1* mRNA. REMSA analysis verified that APUM1/2/3 can specifically bind to the 3’-UTR of *PAP1* mRNA. In addition, *APUM1/2/3* themselves are induced by various stress conditions. Thus, our data suggest that APUM1/2/3-mediated posttranscriptional regulation integrates environmental and developmental cues to precisely modulate anthocyanin biosynthesis, provide a new regulatory layer that fine-tunes anthocyanin production under stress.

## EXPERIMENTAL PROCEDURES

### Plant materials and growth conditions

*Arabidopsis thaliana* Col-0 plants were used as the wild-type for this study. The T-DNA insertion mutants of *apum1* (SALK_100910C), *apum2* (SALK_016283), *apum3* (SALK_076082), *apum4* (SALK_004686C), *apum5* (CS825068) and *apum6* (CS873162) was obtained from the ABRC stock center (https://abrc.osu.edu/) and the homozygous plants were identified by PCR-based genotyping using the primers listed in Table S6. *apum1 apum2 apum3* (abbreviated as *apum1/2/3*) triple mutant was constructed using CRISPR/Cas9 technology. After screening and PCR-based sequencing, two lines of the *apum1/2/3* homozygous triple mutants were obtained for the follow up study.

Seeds were surface sterilized and stratified at 4°C for 3 days, and then germinated on half-strength Murashige and Skoog (MS) medium containing 1% sucrose and 0.8% phytoagar with or without various concentrations of NaCl, mannitol and ABA at 22°C under long-day conditions (16 h light/8 h dark). Seven-day-old seedlings were transplanted to the soil and grown under long-day conditions with a light intensity of 110 μmol photons m^-2^ sec^-1^. The root growth assay was carried out by vertical culture on 1/2 MS medium for certain days. The root phenotype images were scanned and the root length was measured by Image J. The germination rate and greening rate were scored daily in three independent experiments (60–80 seeds per experiment).

### Plasmid construction and plant transformation

The coding DNA sequences (CDS) of *APUM1-APUM6* without the stop codon were cloned into the pENTR SD/D-TOPO entry vector (Invitrogen). For transient subcellular localization analysis, the CDS in pENTR SD/D-TOPO vectors of *APUM1-APUM6* were recombined into the p2GWY7 destination vector (http://gateway.psb.ugent.be/vector/show/p2GWY7/search/index/) and transformed into the Arabidopsis protoplasts according to the method of Wu et al. (2009). Protoplasts were observed and photographed using a laser confocal fluorescence microscope (Olympus, Tokyo, FV 3000). For the construction of *apum1/2/3*, the proper sgRNA sequence (GGATTTGAGGGTTGCTCAG) targeted to the *APUM1*, *APUM2* and *APUM3* was selected using online sgRNA design website (CRISPR-GE http://skl.scau.edu.cn/, CRISPR-P http://cbi.hzau.edu.cn/crispr/). Then, the target sequence was constructed into the AtU6:sgRNA-pYAO:Cas9 high efficiency vector and transformed to the Arabidopsis plant via agrobacterium-mediated transformation.

### RNA isolation and RT-qPCR analysis

Total RNA was extracted using RNAgents Total RNA Isolation System (Promega) and treated with DNase using a DNA-free Kit (Applied Biosystems). The RNA was then reverse transcribed with AMV reverse transcriptase (Promega) according to the manufacturer’s instructions. Gene expression was determined using gene-specific primers by quantitative real-time PCR. Primers used are listed in Table S6. Three biological repeats were analyzed, and each sample was run in triplicate. The data set was normalized using *ACTIN2* as a reference.

### Protein expression, purification and RNA electrophoretic mobility shift assay (REMSA)

The sequences encoding Homology Domain (HD) of APUM1, APUM2 and APUM3 were cloned into the expression vector pGEX-6P-3, and the expression of the proteins were induced using 0.2 mM isopropyl-β-D-thiogalactopyranoside at 16°C overnight in *Escherichia coli* strain Rosetta. The protein was purified as described by Wu et al. (2016) (Wu et al., 2016). The Cy5-labeled probes (P1-P3) were used at 10 nM, and unlabeled competitorswere used at 1 μM. The RNA probes were chemically synthesized, and their 5’-end were labeled by Cy5 (Sangon Biotech (Shanghai) Co., Ltd). The corresponding mutated and non-labeled probes were also chemically synthesized for negative control or competition assays. For REMSA assay, APUM1-HD/APUM2-HD/APUM3-HD proteins and RNA probes were incubated at room temperature for 30 min in the reaction solution (10 mM HEPES pH 7.3, 20 mM KCl, 1 mM MgCl2, 1 mM dithiothreitol, 5% glycerol and 2 μg tRNA). The reactants were analyzed using a native polyacrylamide gel and the Cy5 signals were detected by Azure Biosystems C600. The sequences of the probes were listed in Table S7.

### Transcriptomic analysis

Raw sequencing reads were processed by removing adaptor sequences and low-quality reads. The remaining high-quality reads were aligned to the *Arabidopsis thaliana* reference genome (TAIR10) using HISAT2 (Kim et al., 2015). Gene expression levels were quantified as fragments per kilobase per million mapped reads (FPKM). Differentially expressed genes (DEGs) between samples were identified using DESeq2 (Love et al., 2014), with significance thresholds set at false discovery rate *p* < 0.05 and |log2(fold change)| ≥ 2 (Benjamini-Hochberg correction). Functional enrichment analysis of DEGs was performed using agriGO v2.0 (https://metascape.org/gp/index.html#/citations) to identify overrepresented Gene Ontology (GO) terms.

### Anthocyanin induction and measurement

For anthocyanin induction, the wild type and *apum1/2/3* were surface sterilized and stratified at 4°C for 5 days. The seeds were germinated in sucrose-free half-strength MS solution for 4 days on a shaker at 120 rpm. Four-day-old seedlings were then transferred to the 0% and 5% sucrose-containing half-strength MS solution for 2 days. Then, seedlings were harvested for anthocyanin measurement using the method described previously by Rabino and Mancinelli (1986) (Rabino and Mancinelli, 1986).

### Accession Numbers

Sequence data from this article can be found in the TAIR data libraries under the following accession numbers: APUM1 (At2g29200), APUM2 (At2g29190), APUM3 (At2g29140), APUM4 (At3g10360), APUM5 (At3g20250), APUM6 (At4g25880).

## Acknowledgments

We thank Mengjiao Chen from Shanghai Normal University for helping upload the RNA seq data.

## Funding

This work was supported by funds from the Shanghai Natural Science Foundation (21ZR1447100) and the National Science Foundation of China (32100191); Shanghai Natural Science foundation (21ZR1447100) and Innovation Program of Shanghai Municipal Education Commission (2021-01-07-00-02-E00117).

## Author Contributions

Wu W and Huang J designed the study. Li D, Xu W, Lin D, Chen T and Chen X made transgenic plants and mutants. Li D, Chen T, Chen X and Lin D perform the physiological experiment, Li D, Xu W, Guo W analyzed the anthocyanin content and related gene expression. Wu W and Li D perform the RNA binding assay. Long Z, Tu X, Xu X, Wu W, Huang J and Chen X prepared and analyzed the RNA-seq results. Wu W and Huang J wrote the manuscript. All authors read and approved the manuscript.

## Declaration of interests

The authors declare no competing interests.

## Supplementary Figure Legends

**Supplemental Figure 1.** Identification of the T-DNA insertion mutants of *apum1*, *apum2*, *apum3*, *apum4*, *apum5* and *apum6*. **A)** T-DNA insertion positions were shown in *APUM* genes. **B)** RT-PCR analysis of expression of APUMs in their corresponding T-DNA insertion mutants.

**Supplemental Figure 2.** Triple *apum1/2/3* mutants constructed by CRISPR-Cas9 technique. **A)** The sgRNA target sites location of *APUM1*, *APUM2* and *APUM3*. **B)** Nucleotide changes of *APUM1*, *APUM2* and *APUM3* in *apum1/2/3*. **C)** Amino acid changes of apum1, apum2 and apum3 in *apum1/2/3*.

**Supplemental Figure 3.** REMSA analysis of the binding of APUM1-HD, APUM2-HD and APUM3-HD to the NRE sequence. P1 represents the Cy5-labelled NRE probe. Bound represents the band of protein binding to the probes. Free represents the unbound probe.

**Supplemental Table 1.** Differentially expressed genes (DEGs) significantly mis-regulated in *apum1/2/3* compared to WT (*p* < 0.05).

**Supplemental Table 2.** DEGs upregulated more than log2(fold change) ≥ 2 in *apum1/2/3* compared to WT.

**Supplemental Table 3.** DEGs downregulated more than log2(fold change) ≥ 2 in *apum1/2/3* compared to WT.

**Supplemental Table 4.** Genes upregulated more than log2(fold change) 1.5-fold in *apum1/2/3* contain at least one predicted APUM-binding sites that are searched with the NRE motif 5’-UGUAUAUA-3’ or its variants with one-altered nucleotide except in position 2 in the 3’-UTR.

**Supplemental Table 5.** Prediction of genome-wide APUM-binding sites in *Arabidopsis* mRNA 3’-UTR. The core sequence 5’-UGUGUAUA-3’ or its variants with one-altered nucleotide except at position 3 were searched in the 3’-UTR of the whole transcriptome. “Yes” means upregulated genes in *apum1/2/3*.

**Supplemental Table 6.** Primers used in this study.

**Supplemental Table 7.** The sequences of the probes used in REMSA.

## Notes

### Competing Interest Statement

The authors have declared no competing interest.

