## Supplementary figures and images for "*Arabidopsis* Group I Pumilio RNA-binding factors are vital for embryo development and balancing between growth and stress resistance"

### Supplemental Figure

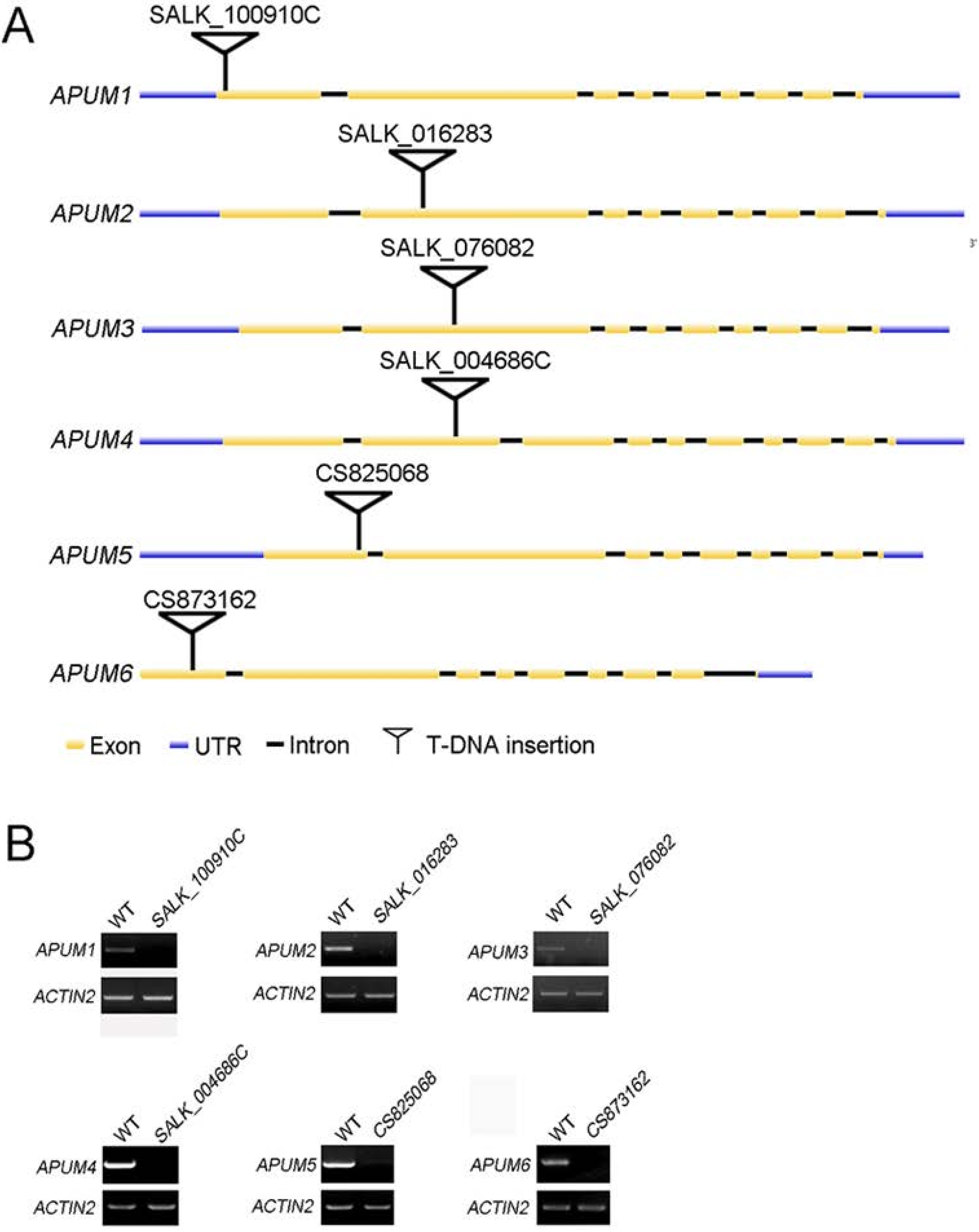

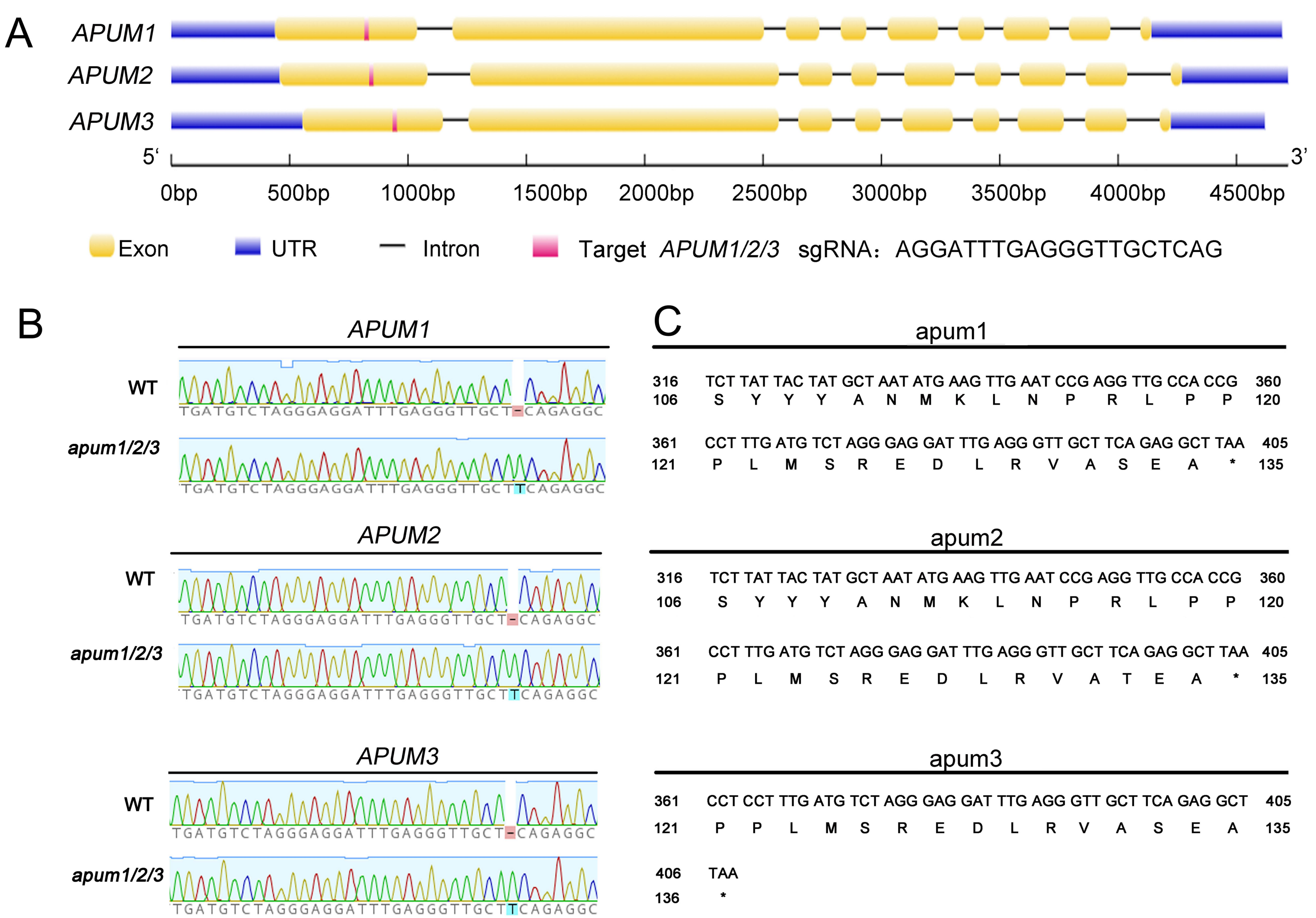

P1: CY5-NRE

CY5-CCAGAAUUGUAUAUAUUCG

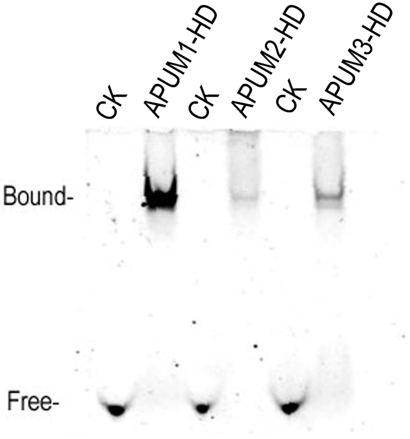
