## Supplemental Table 6 for "*Arabidopsis* Group I Pumilio RNA-binding factors are vital for embryo development and balancing between growth and stress resistance"

Table S6 Primers used in this study.

| Primers name | Primer sequence (5’ to 3’) | Objective |
| --- | --- | --- |
| *APUM1-F* | TCCCATTCAGAAGCAGAAGCACCAG | *apum1* (SALK_100910C) mutant identification |
| *APUM1-R* | AGATCTCAAAACAGTTCCAACGAG |  |
| *APUM2-F* | AGAGGAAGCAGCATGAGTTTG | *apum2* (SALK_016283) mutant identification |
| *APUM2-R* | ACCCATAAAGTTCACGTCCATG |  |
| *APUM3-F* | GTGGGGCTTCTTTTCTGGAA | *apum3* (SALK_076082) mutant identification |
| *APUM3-R* | TGCACGAAGCAGTTCCCCTA |  |
| *APUM4-F* | ATTTGACAGCCCATTAGCATTC | *apum4* (SALK_004686C) mutant identification |
| *APUM4-R* | AGGTATTCGTTAGCCGTATCTCC |  |
| *APUM5-F* | AATAGAAGTATGACGACTACGCAGA | *apum5* (CS825068) mutant identification |
| *APUM5-R* | ACGATGAAGGTGTGTATTGTGG |  |
| *APUM6-F* | CACCATGGCAACTGAGAATCCTATTAG | *apum6* (CS873162) mutant identification |
| *APUM6-R* | GCTGGATTCCATAGTAATTAACAC |  |
| *Lba1* | TGGTTCACGTAGTGGGCCATCG | T-DNA insertion identification |
| *Lb3* | TAGCATCTGAATTTCATAACCAATCTCGATACAC |  |
| *APUM123-C-F* | GATTGGGATTTGAGGGTTGCTCAG | AtU6:APUM123sgRNA-pYAO:Cas9 construction |
| *APUM123-C-R* | AAACCTGAGCAACCCTCAAATCCC |  |
| *APUM1-D-F* | GTGGGGCTTCTTTTCTGGAA | *APUM1* target sit amplification |
| *APUM1-D-R* | CTATACACAACACATACACAC |  |
| *APUM2-D-F* | ATGTTTCTTTTCCTGAAGAGTTTAGTG | *APUM2* target sit amplification |
| *APUM2-D-R* | AGAAAGAAGCTAACAAGAAGCGTTGTC |  |
| *APUM3-D-F* | TGGTGTGAGTATAAACTATTG | *APUM3* target sit amplification |
| *APUM3-D-R* | CTATAGACAACCAAAAGAAGTA |  |
| *APUM1-RT-F* | AGAAGACCGATGCATAGAGGT | *APUM1* RNA level detection |
| *APUM1-RT-R* | AGCACCAGTTTTCTCACCTTC |  |
| *APUM2-RT-F* | AGAGGAAGCAGCATGAGTTTG | *APUM2* RNA level detection |
| *APUM2-RT-R* | CATCATTATAGGTAGATCCTTGTGC |  |
| *APUM3-RT-F* | GTGGGGCTTCTTTTCTGGAA | *APUM3* RNA level detection |
| *APUM3-RT-R* | TCATTTGTGAATTCCTGCACGAAG |  |
| *APUM4-RT-F* | ATTTGACAGCCCATTAGCATTC | *APUM4* RNA level detection |
| *APUM4-RT-R* | AGGTATTCGTTAGCCGTATCTCC |  |
| *APUM5-RT-F* | AATAGAAGTATGACGACTACGCAGA | *APUM5* RNA level detection |
| *APUM5-RT-R* | ACGATGAAGGTGTGTATTGTGG |  |
| *APUM6-RT-F* | ATGATGATGAAGGACCAGTATG | *APUM6* RNA level detection |
| *APUM6-RT-R* | TCATCTCCTCAATTCTTGGTTTT |  |
| *ACTIN2-RT-F* | TCTTCTTCCGCTCTTTCTTTCC | *Actin2* RNA level detection |
| *ACTIN2-RT-R* | TCTTACAATTTCCCGCTCTGC |  |
| *qPUM1-F* | TATGCCACCGGGTTTTGAAGG | *APUM1* RNA level quantification |
| *qPUM1-R* | TATGCCACCGGGTTTTGAAGG |  |
| *qPUM2-F* | TATGCCACCGGGTTTTGAAGG | *APUM2* RNA level quantification |
| *qPUM2-R* | TATGCCACCGGGTTTTGAAGG |  |
| *qPUM3-F* | TATGCCACCGGGTTTTGAAGG | *APUM3* RNA level quantification |
| *qPUM3-R* | TATGCCACCGGGTTTTGAAGG |  |
| *qPUM4-F* | TGAGCACTATTCCGCTGAAGGC | *APUM4* RNA level quantification |
| *qPUM4-R* | ACTGGAGAGTGAAGCTCAGAAGC |  |
| *qPUM5-F* | GGTCAAGTCGCCACTCTTTC | *APUM5* RNA level quantification |
| *qPUM5-R* | CTCCAGGACATGCTGAGTGA |  |
| *qPUM6-F* | CATTGTTCAGCCGAGTGAGA | *APUM6* RNA level quantification |
| *qPUM6-R* | TGAGTCAGCCCACTGTCTTG |  |
| *qACTIN2-F* | CACTTGCACCAAGCAGCATGAAGA | *ACTIN2* RNA level quantification |
| *qACTIN2-R* | AATGGAACCACCGATCCAGACACT |  |
