## Supplemental Table 7 for "*Arabidopsis* Group I Pumilio RNA-binding factors are vital for embryo development and balancing between growth and stress resistance"

Table S7 The sequences of the probes used in REMSA.

| name | Sequence (5’-3’) |
| --- | --- |
| P1:Cy5-NRE | (CY5)CCAGAAUUGUAUAUAUUCG |
| P2:Cy5-mNRE | (CY5)CCAGAAUUUUAUAUAUUCG |
| C1:NRE | CCAGAAUUGUAUAUAUUCG |
| P3:Cy5-PAP1 | (CY5)GCGUUUGUGUAUAGGUUUGC |
| C2:PAP1 | GCGUUUGUGUAUAGGUUUGC |
| C3:m3PAP1 | GCGUUUUUUUAUAUAUUUGC |
| C4:m4PAP1 | GCGUUUAAAAAAAAAUUUGC |
| C5:m5PAP1 | GCGUUUGCGCGCGGGUUUGC |
| mC1 | GCGUUCGUGUAUAGGUUUGC |
| mC2 | GCGUUUUUGUAUAGGUUUGC |
| mC3 | GCGUUUGCGUAUAGGUUUGC |
| mC4 | GCGUUUGUUUAUAGGUUUGC |
| mC5 | GCGUUUGUGCAUAGGUUUGC |
| mC6 | GCGUUUGUGUGUAGGUUUGC |
| mC7 | GCGUUUGUGUACAGGUUUGC |
| mC8 | GCGUUUGUGUAUGGGUUUGC |
| mC1+4 | GCGUUCGUUUAUAGGUUUGC |
| mC2+4 | GCGUUUUUUUAUAGGUUUGC |
| mC5678 | GCGUUUGUGCGCGGGUUUGC |
